# Communication between wolf and domestic dog revealed from experimental scent marking

**DOI:** 10.1101/2023.06.04.543479

**Authors:** Kinga M. Stępniak, Tom A. Diserens, Maciej Szewczyk, Robert W. Mysłajek, Dries P.J. Kuijper

**Affiliations:** Department of Ecology, Institute of Functional Biology and Ecology, Faculty of Biology, University of Warsaw, Biological and Chemical Research Centre, Żwirki i Wigury 101, 02-089 Warszawa, Poland; Mammal Research Institute Polish Academy of Sciences, Stoczek 1, 17-230 Białowieża, Poland; Department of Vertebrate Ecology and Zoology, Faculty of Biology, University of Gdańsk, Wita Stwosza 59, 80-308 Gdańsk, Poland

**Keywords:** chemical communication, intraspecific interactions, conservation, territory defense

## Abstract

The European grey wolf (*Canis lupus*) and the domestic dog (*Canis lupus familiaris*) share not only a common origin but also many similarities in their behavior. Due to the implementation of protection wolves repopulated large parts of Europe. The increase of wolves in human-dominated landscapes also leads to a growing potential for interactions between wolves and domestic dogs. However, these interactions between wolves and dogs are still poorly understood. Scent marking is one of the main forms of communication in canids, as is crucial for territorial marking, synchronization of reproduction, establishment of hierarchies in groups, and formation of new breeding pairs. We hypothesized that the presence of scent markings by domestic dogs in wolf territories elicits a behavioral response of wolves and therefore may interfere with natural wolf behavior. To test this, we experimentally scent-marked objects within known wolf home-ranges in the Kampinos National Park (Poland) to simulate the presence of “unknown dogs” (dog urine from outside the area) and water as a control. To test whether and how the behavioral response differs between domestic dogs and wolves we additionally created scent marks of “unknown wolves” (wolf urine from outside the area). By means of camera traps we studied the behavioral repones of local wolf families exposed simultaneously to all three scent stimuli. Our study showed that wolves (breeding pair) reacted to scent marking from “unknown dog” in 16% of cases, while an average juvenile reacted in 27% of cases. In 33% of cases, the breeding pair overmarked stimuli from an “unknown dog” and in 27% of cases mark them by ground scratching. Wolves spend significantly more time exploring and sniffing scent marks of “unknown wolves” than “unknown dogs”.

Our result indicates that domestic dog scent marks trigger a behavioral response in wild wolves showing that it does affect their behavior. The longer time that wolves spend on responding to wolf scent marks compared to dog scent marks indicates they can distinguish between wolf and dog scent marks, but especially inexperienced juveniles spend much time exploring dog scent marks. This suggests that the increasing occurrence of dogs inside wolf territories could affect and potentially disturb the scent-marking behavior of wolves.

## Introduction

Due to the implementation of strict protection wolf (*Canis lupus)* populations increased their range and numbers across Europe (Chapron et al. 2014, Nowak and Mysłajek 2016). The recovery of wolves in human-dominated landscapes often brings them into contact with ‘humans’ best friend’ the domestic dogs (*C. l. familiaris*). In many places around the world, wolves and dogs live close to each other, or even share the same landscape (Wierzbowska et al. 2016, Mysłajek et al 2018, Salvatori et al 2020, Mysłajek et al 2022).

This creates a fascinating situation when wild and domesticated forms of this same species have a possibility to meet again. On the other hand, it also raises a lot of questions, mostly about the consequences of their interactions with each other.

Wolves and dogs can interact in several ways. They are closely related, share identical karyotypes, can interbreed, and produce fertile offspring (Salvatori et al. 2020). In addition to hybridization, dogs can be competitors to wolves for similar food resources (Hughes and Macdonald 2013). Therefore, in places where the dog presence overlaps with wolves’ territories, dogs can compete with the wolf, limiting its access to resources. Moreover, dogs and wolves share many parasites and diseases (Lescureux and Linnell 2014). Among the numerous pathogens affecting dogs, only a few are believed to be of concern for wolf conservation, but their presence in the wild can have severe consequences (Knobel et al. 2014). Thus, there is a growing concern among conservationists about the negative impacts of dogs on wolf behavior and ecology (Lescureux and Linnell 2014).

While competitive interactions and parasites were widely studied, the behavioral response of wolves to dog presence within their territories has generally been overlooked. Dogs and wolves share similar behaviour, even though in some breeds it has been distorted by selective breeding or has lost its function during domestication. Scent marking is one of the major forms of communication for canids, including wolves (Peters and Mech 1975, Dunbar 1977, Allen et al. 1999, Pal 2003, Zub et al., 2003, Stępniak et al. 2020). Similar to wolves, dogs also use scent markings to communicate with conspecifics (Scott and Fuller 1965, Pal 2003). Scent marks contain a rich source of information that largely affects the behavior of both dogs and wolves. In wolves, it entails information essential for marking territories, pair formation, reproduction, the establishment of hierarchies in groups, and formation of new breeding pairs (Asa et al. 1985, Paquet and Fuller 1990, Paquet 1991, Vila et al. 1994).

Despite the general lack of information, it is clear that also for dogs the use of scent marking is a crucial component of their communication in a broadly similar way as for wild wolves. Scent marking in canids include feces, urine, and interdigital gland secretions left during scratching the ground (Harrington and Asa 2003). Since the production of pheromones is closely related to the level of hormones, it carries a range of information (e.g., regarding sex, health or position in the group) of the individual that left them (Zala et al. 2004, Wyatt, 2014).

The cost of scent marking is much lower than the cost of attacking or being attacked by an intruder, therefore scent marking is an economic strategy to protect the wolf’s territory and its resources against potential intruders (Gosling 1982, 1986, Gosling and Roberts 2001). The intensity of scent marking varies seasonally and is influenced by the status of the individual (Peters and Mech 1975, Asa et al. 1990). Especially after successful reproduction, wolves are increasing the number of scent markings left near to the core areas of their territories (Zub et al. 2003, Llaneza et al., 2014). To enhance information transfer, wolves often leave their scent marks in conspicuous places like forest roads intersections, mostly on conspicuous surfaces and objects (Zub et al. 2003, Stępniak et al. 2020), similarly to free-living dogs (Cafazzo et al. 2012).

As wolves and dogs are wild and domestic versions of the same species and scent marking is a crucial way of communication for both, it is likely that they interact via scent marking when they occur in the same area. However, what information they transfer and how this affects wolf behaviour is yet unknown. The presence of dogs could be perceived by wolves as indicators of risk from humans (Suraci et al. 2019). Dogs can also be recognized as competitors (Hughes and Macdonald 2013). In this case, wolves should rather avoid dogs or their scent marks, and the presence of dogs could exacerbate the human fear effects.

Alternatively, dogs may be perceived as potential reproductive mates (Lescureux and Linnell 2014), which could lead to wolves being attracted to dogs or their scent marks. Whether and to what extent wolves and dogs communicate with each other via scent marks and how this may alter wolf behaviour is therefore an important knowledge gap in the light of the increasing presence of wolves in human- and dog-dominated landscapes in Europe.

To test whether and to what extent wolves react to scent marks by dogs and how their response differs from reactions to wolves, we experimentally scent-marked locations to simulate the presence of “unknown dogs” (dog urine from outside the area), “unknown wolves” (wolf urine from outside the area) and as well as water as a control. By means of camera traps we studied the behaviorial reponse of wolf families living in a by humans and dogs intensively visited area (Kampinos National Park).

## Methods

### Study area

Kampinos National Park (hereinafter KPN) was established in 1959. Its total area is 384.7 km^2^, and the buffer zone around the Park covers additional 377.6 km^2^. The area is also protected as the Natura 2000 site “Puszcza Kampinoska” (PLC140001), which covers 376.4 km^2^. In 2000, the KPN and its buffer zone were assigned UNESCO-MaB (Man and Biosphere) program as a Biosphere Reserve. The landscape of the park was shaped in the post-glacial period, which resulted in the formation of marsh and dune areas. KPN covers the area of the Kampinos Forest in the Vistula proglacial stream valley, in the NW part of Warsaw, between the left bank of the Vistula and Bzura rivers (fig.1A). Forests cover over 73% of the park’s area, with the Scots pine (*Pinus sylvestris*) being the dominant species (70%), followed by the black alder (*Alnus glutinosa*) and oaks (*Quercus* ssp.), 12.5% and 10.3%, respectively.

**Figure 1.**
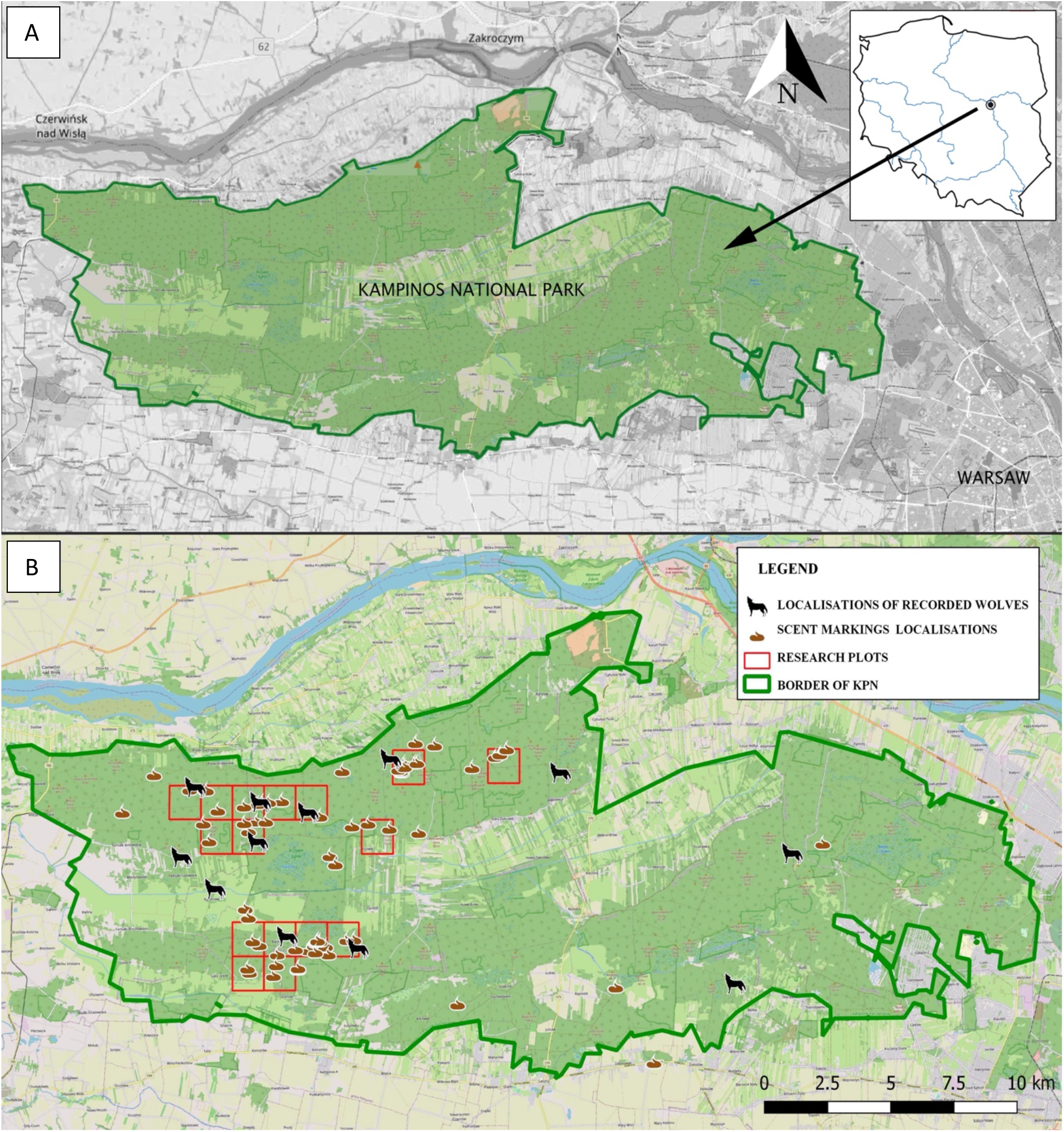
A/ Study area, Kampinos National Park. B/Distribution of research areas, scent markings and places where wolves were recorded.

The ungulate community consists of the roe deer (*Capreolus capreolus*) (78%), red deer (*Cervus elaphus*) (9%), moose (*Alces alces*) (8%), and wild boar (*Sus scrofa*) (5%) (Central Statistical Office 2019). In 1994, the KPN started reintroduction of the Eurasian lynx (*Lynx lynx*) (Böer et al. 2000) and the presence of this species is still confirmed (Mysłajek et al. 2019). After wolves were put under protection in 1998 (Nowak and Mysłajek 2017), this species increased its range and recolonized Kampinos National Park. The first confirmed reproduction in KPN happened in 2015. During the study period, two wolf families live in KPN, and their total number was estimated at approx. 12 individuals (K.M. Stępniak, M. Szewczyk, I. Kwiatkowska, R.W. Mysłajek, S. Nowak, unpublished data).

The Kampinos National Park, due to its location right next to Warsaw, is very popular place for recreation for residents of the capital. All of the KPN’s borders are surrounded by human settlements, and some villages are located within the park’s boundaries. This makes much of the national park under heavy anthropogenic pressure. Moreover, because dogs’ presence in protected areas is correlated with human-related factors (Soto and Palomeres 2014), these canids are highly abundant in KPN.

Due to art. 30 (Journal of Laws 2022.0.672 i.j. - Act of 28 September 1991 on forests), dogs should enter the forests only on a leash. Regulations of the Kampinos National Park prohibit the introduction of dogs into areas under strict and active protection, except for assistance dogs (order no. 15/2020 Director of the Kampinos National Park dated 31.07.2020 on the improvement of the Kampinos National Park).

### Selection of sample plots

We located our sample plots in areas with the highest chance of wolf presence based on the analysis of data collected during two previous research projects in KPN:

1. data from the camera traps installed in dense networks in 68 fixed locations in the KPN (see Suppl. Material. S1). Data were collected between 03.05.2018 and 28.06.2018, which gave a total of 2366 camera trapping days (unpubl. results Mammal Research Institute PAS). Camera traps were deployed along forest roads, on trees at height of up to 1m, in places where there was at least 20m visibility (following Bubnicki et al. 2019),
2. Data from four years (2016-2020) of wolf winter tracking in KPN. During each tracking, we collected non-invasive genetic samples (i.e., scats and urine). Coordinates of each sample were marked on a hand-held GPS receiver (Garmin GPSMAP® 64s) and then entered into the database (following Szewczyk et al. 2019, Stępniak and Kwiatkowska unpublished data).

Using QGIS program (QGIS.org, 2021) we plotted a regular grid with cells of 1.25 km^2^ on the map of the surface of the KPN together with all locations with wolf presence based on these two previous studies. As areas of high wolf activity, we selected squares where two or more wolf observations were present (fig.1B). Within these cells we selected plots for the experiment during field work. While moving along areas of high wolf activity, we choose intersections (Barja et al. 2004, Stępniak et al. 2020) with scent markings indicating recent wolf presence and with sufficient trees for camera trap installation and with full visibility of the experimental plots.

All experimental plots were placed in the territory of one family group living in KPN (fig. 1B). Moreover, during an analysis of data collected during the experiment, we observed reactions to stimuli performed by both wolves’ groups.

### Study design

We created three treatments by adding the following scent marks: (1) dog urine and (2) wolf urine, and (3) water as a control. We replicated this set of three treatments 29 times. At each treatment plot, we added 10ml of treatment at a height not higher than 0.5 m from the ground surface. To prevent contamination between individuals’ scents, we used one new syringe for each scent stimulus. Moreover, to reduce contamination with human scent while applying the treatments, we wore nitrile gloves. To mimic natural scent marking, all three types of scent stimuli were placed on distinctive elements of the environment (trees, clumps of grass, tree trunks, bushes, etc.). At each location (i.e. forest crossroads) we placed scent stimuli in random order to prevent those wild wolves would be presented with one stimulus first. Each week, urine from a new individual wolf and dog (“unknown wolf” and “unknown dog” see details under ‘*Wolf and dog urine collection’*) was used to give the appearance of new individuals appearing in the wolf territory.

To record wolf behavioural response to scent stimulus, we installed a camera trap (Ltl Acorn 5310 MG or BUSHNELL Trophy cam HD) on the tree at the height of up to 0.5 m from the ground with an unobscured camera trap view aimed at the exact location of our applied scent stimuli, at 8-10 m. The camera traps were set to record films with a length of 60 seconds from the moment of triggering. The interval between subsequent recordings was set at 1 second. Camera traps were installed for 7 days to minimize the effects of aging of scent stimuli. When wolves were recorded at a location after setting up the treatments, we moved to a new location. When no wolves were recorded, we continued at the same location and refreshed scent marks at the beginning of each week. The GPS coordinates of each treatment plot were recorded and to facilitate video analyses of the wolves’ response toward the treatments we temporarily marked each scent plot at the start of the experiment with a number written on a piece of paper and photographed it. Afterward, the piece of paper was removed.

As potentially the response to a new dog scent mark is triggered or modified by the presence of a new wolf scent mark, we established four additional locations with treatment plots. At these locations, only two treatments were placed, urine from an unknown dog and water. This allowed us to test whether the response towards a new dog is different with and without the presence of a new wolf scent mark. The selection process of these plots was similar to the selection of experimental plots on which all three types of stimuli were placed.

### Wolf and dog urine collection

In the experiment, we used urine from five wolves (two females and three males) and from six dogs (three females and three males). Wolf urine was collected from the wolves at the Wolf Science Center of the Veterinary University of Vienna. The wolves kept there are under constant veterinary care, vaccinated, and free of internal and external parasites. Urine was collected by wolf trainers directly from the wolf into sterile plastic containers.

We collected domestic dog urine from healthy, vaccinated, and dewormed individuals living in private homes, directly from the dog into sterile plastic containers. Urine was stored in freezers for 18 months. The use of frozen urine is an accepted and used research method (Lisberg and Snowdon, 2009). The permission for the study was granted by the Director of the Kampinos National Park. According to the Polish Law on Use of Animals in Scientific Experiments for the study which does not involve direct interaction with animals’ permission from the ethical commission is not required.

### Video classification

The classification of recordings was performed in the “Trapper” software (Bubnicki et al. 2016). Details of classifiers are shown in Table 1.

**Table 1.**

| Classifier name | Classifier options |
| --- | --- |
| the type of recording | observation, blank, camera trap inspection, recording error |
| the number of individuals visible on the recording |  |
| species | wolf, dog with a human on a leash, dog with a human but not on the leash, dog without a human present |
| status (classification only for wolves) | breeding female, breeding male, juvenile, unspecified |
| reaction to a scent stimulus? | yes, no |
| type of stimulus to which the reaction was recorded | wolf, dog, water |
| type of reaction to the scent stimulus | sniffing, overmarking, ground scratching |
| reaction time | counted in seconds |
| time of presence on the recording | counted in seconds |
| was the stimulus overmarked by another animal? | yes, no |
| the species of animal that overmarked the stimulus | fox, dog, badger, wolf |

Differentiation by the reproductive status and sex of recorded wolves was made based on detailed observation of the wolf visible in the recording (following Goodman et al. 2002). The following factors determined the categorization of a wolf into a given group (breeding pair or juveniles): the size of the wolf relative to other members of the pack, posture, tail position, and behavior. The breeding pair usually were clearly larger than the subadults, carried their tail high as a sign of social dominance, their body posture was dominant. Secondary sex characteristics and posture during stimulus overmarking were used to discriminate between sexes in the breeding pair. Males overmarked by lifting a raised leg urination and female by lifting a bent hind leg and gently squatting (Peters and Mech, 1975)

We measure the duration of each reaction separately. The behavior of each individual wolf and each individual dog present on the recording was classified.

### Statistical analysis

The analysis of wolf and dog behavior was performed in Microsoft Excel and the visualization of the results was performed in SigmaPlot. Statistical significance analysis of the obtained results was performed in the R program (R Core Team 2021) using two-factor ANOVA test and Tukey-HSD test. Time-activity analysis was performed in R program using the extension “camtrapR” (Niedballa et al. 2016).

## Results

### Behavioral response of all wolves towards scent marks

When wolves showed a response, they spent the most time sniffing the stimuli an average of 52% (SE ± 0.040) of their time they were visible on recordings. They spend on average 8.4%, (SE ± 0.041) of the total time on recordings on unknown dog stimuli (*F*_2,36_= 12.05; p < 0.001). However, wolves concentrated mainly on sniffing unknown wolf stimuli, by spending 43.6% (SE ± 0.067) of the total time during which they were seen on the recordings. The second behavioral response with the longest duration to scent marks was scratching the ground, which took 4.2% (SE ± 0.008) of the total time during which they were seen on the recording. Wolves spend responding to the unknown dog stimulus (average 1.7% ± SE 0.017) and to the unknown wolf stimulus (average 2.5% SE ± 0.018) of responding time. These differences were not statistically significant (*F*_2,36_= 1.409; p > 0.05). Wolves spent the least of total time (mean 0.9%, SE ± 0.002) during which they were visible on the recording, on overmarking stimuli. Unknown dog stimuli were overmarked an average of 0.5% of the time (SE ± 0.005) and unknown wolf stimuli were overmarked an average of 0.4% of the time (SE ± 0.003).

### Behavioral response of breeding pair and juvenile wolves

The response of the breeding pair and juveniles to scent marks differed in terms of the time spent on it and the number of responses The only reaction presented by juveniles was sniffing, however, the duration of sniffing of unknown dogs and unknown wolf scent marks differed statistically (*F*_2,36_ = 9.278; p> 0.001). The juveniles together spent 441 seconds sniffing the scent marks, which constituted an average of 53% (SE ± 0.066) of time when they were recorded. They spent more time sniffing stimuli from the unknown wolf than stimuli from the unknown dog (40%; ± 0.074, 13%; ± 0.057, respectively). The remaining time (43%; ± 0.0388) was devoted by juveniles to other behaviors not aimed toward the scent marks (table 2A, figure 2A).

**Table 2A.** Comparison of the effect of scent stimuli on the behavior of breeding pairs and offspring in Kampinos National Park, using the Tukey-HSD test. The asterisk symbol indicates statistical differences between the compared stimuli.

| Individuals | Behavior | Scent stimuli | 95% confidence level |  | Level of statistical significance |
| --- | --- | --- | --- | --- | --- |
|  |  |  | lower | upper |  |
| Breeding pair | sniffing | NW - ND | -0.1006761 | 20.100676 | 0.0527940 * |
|  |  | Water - ND | -11.177599 | 9.023753 | 0.9632908 |
|  |  | Water - NW | -21.177599 | -0.976247 | 0.0289758 * |
|  | overmarking | NW - ND | -1.153465 | 2.0765418 | 0.7658653 |
|  |  | Water - ND | -2.076542 | 1.1534649 | 0.7658653 |
|  |  | Water - NW | -2.538080 | 0.6919264 | 0.3530720 |
|  | ground scratching | NW - ND | -1.153465 | 2.0765418 | 0.7658653 |
|  |  | Water - ND | -2.076542 | 1.1534649 | 0.7658653 |
|  |  | Water - NW | -2.538080 | 0.6919264 | 0.3530720 |
| Juveniles | sniffing | NW - ND | 5.858268 | 40.14173 | 0.0063829 *** |
|  |  | Water - ND | -22.603270 | 11.68019 | 0.7182921 |
|  |  | Water - NW | -45.603270 | -11.31981 | 0.0007272 *** |

**Tabela 2B.**
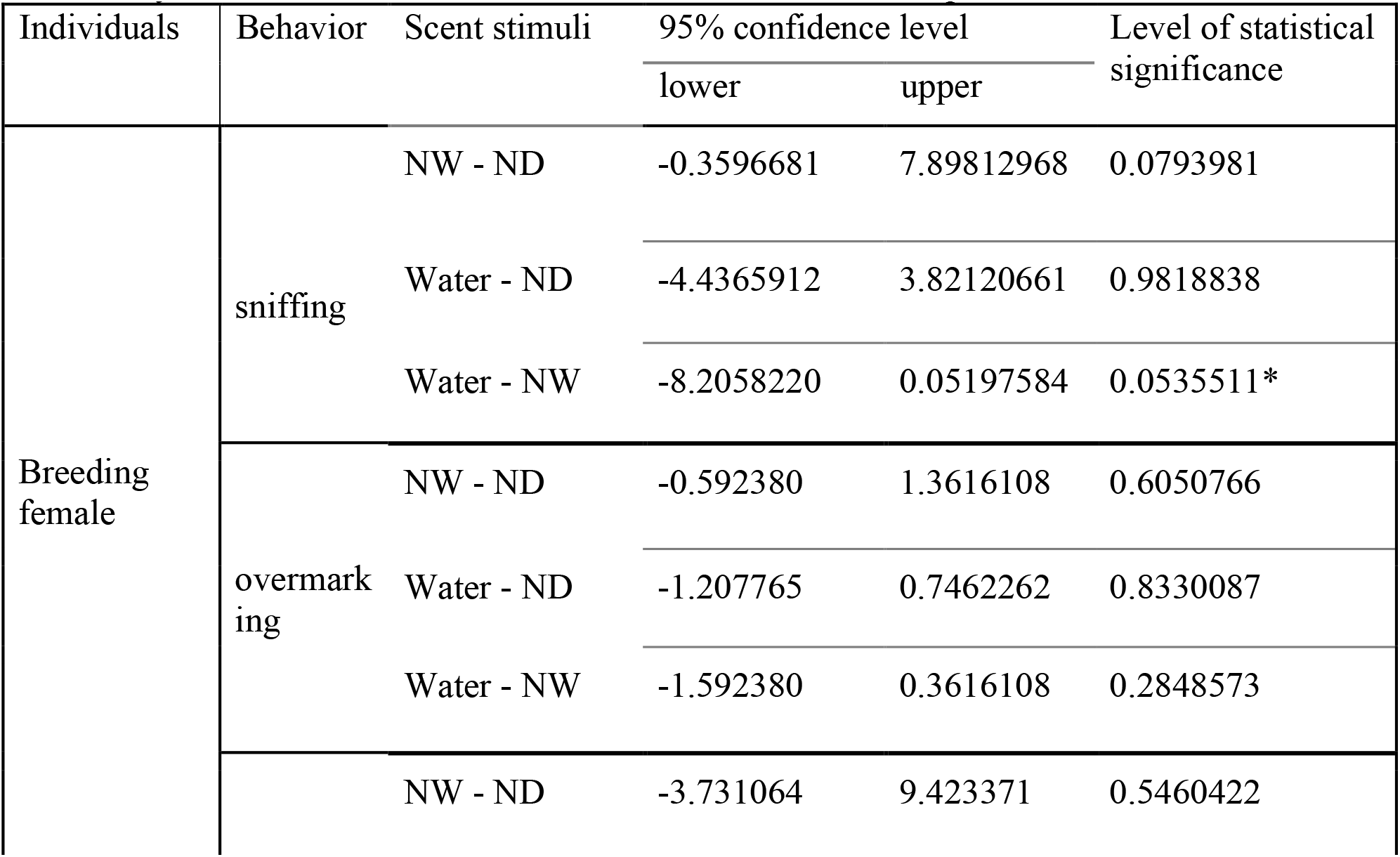
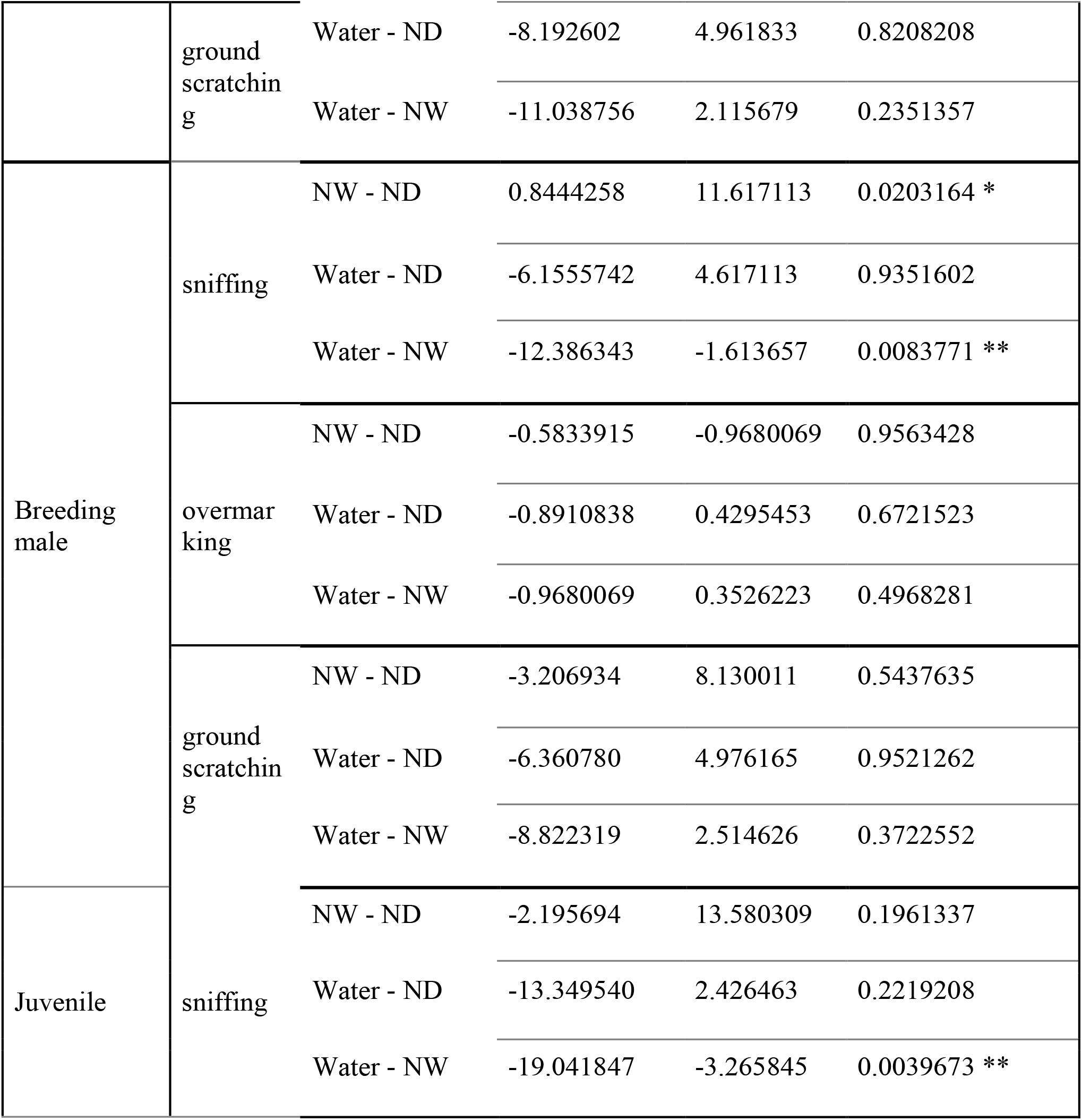
Comparison of the effect of scent stimuli on the behavior of breeding female, breeding male and juvenile in Kampinos National Park, using the Tukey-HSD test. The asterisk symbol indicates statistical differences between the compared stimuli.

**Figure 2A.**
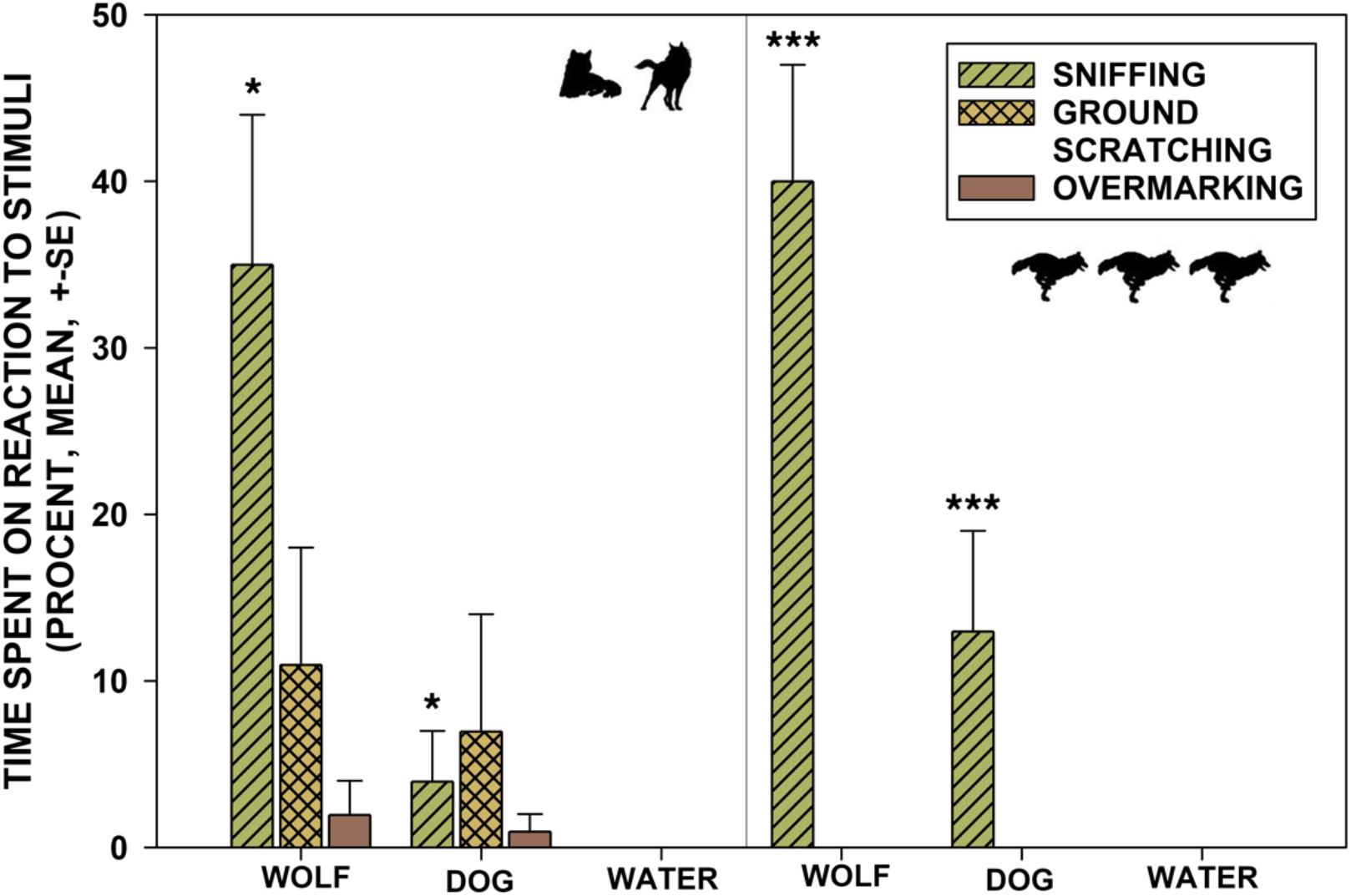
Percentage response length (mean, SE) of breeding pair (left) and juveniles (right) to stimuli from NW, ND and control stimulus (water) in the Kampinos National Park. The asterisk (*) indicates statistically significant differences between the stimuli.

**Figure 2B.**
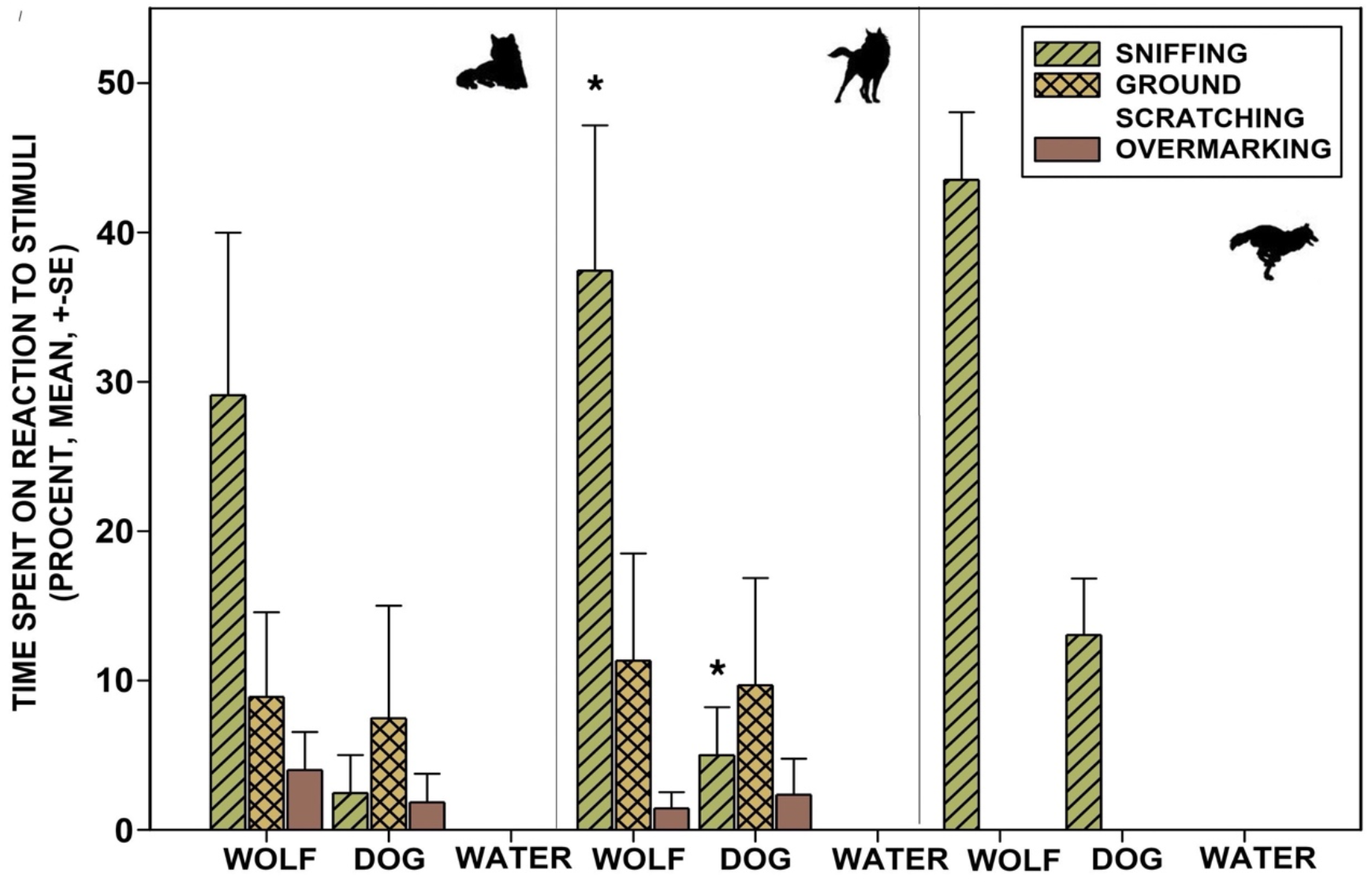
Percentage response length (mean, SE) of a breeding female (left), a breeding male (center) and a juvenile (right) to stimuli from NW, ND, and to a control stimulus (water) in the Kampinos National Park. The asterisk (*) indicates statistically significant differences between the stimuli.ń

The breeding pair spent 255 seconds reacting to all scent marks, which was 59.4% (SE ± 0.02) of time spent on the recording. The most time (mean 38.3%; SE ± 0.048) the breeding pair spent on sniffing the unknown wolf scent mark was sniffed longer than the unknown dog scent mark (mean 34.8; SE ± 0.089, mean 3.5; SE ± 0.026, respectively). Scratching the ground (mean 18.1%; SE ± 0.033) was the second time-consuming reaction to stimuli, reaction to unknown wolf stimuli was stronger than that to unknown dog stimuli (mean 11.1; SE ± 0.071, mean 7.0%; SE ± 0.07, respectively). The shortest time the breeding pair spent on overmarking, mean = 3% (SE ± 0.006) of the entire time when the wolves were visible on the recording (unknown dog mean time 2.1%; SE ± 0.015 unknown wolf, mean time 0.9%; SE ± 0.008). The difference between the time spent sniffing an unknown dog scent mark and the time spent sniffing an unknown wolf scent mark is statistically significant (F = 4.37; p <0.05). The difference between the time spent on overmarking and scratching the ground, between the reactions to unknown dog stimuli and unknown wolf stimuli is not statistically significant (*F*_2,36_= 0.975; p> 0.05 and *F*_2,36_= 1.409; p> 0.05, respectively) (table 2A, figure 2A).

### Behavioral response of male, female and juvenile wolves

The breeding male spent more reacting to scent marks than the other behaviors not aimed at the scent marks (66% and 34% respectively). He spent the main part of this time sniffing scent marks (39%, average 30%, SE ± 0.085), then scratching the ground (24%, average 17%, SE ± 0.077). The shortest part of the time during which a male was visible on the recordings was spent on overmarking stimuli (3%, mean 3%, SE ± 0.017). The difference between the time that male spent on sniffing unknown dog and the time that male spent on sniffing unknown wolf stimuli is statistically significant (*F*_2,36_= 6.069; p <0.01). In the case of overmarking and scratching the ground, the time differences between the reactions to the unknown dog and unknown wolf stimuli are not statistically significant (respectively, *F*_2,36_ = 1.21; p> 0.05 and *F*_2,36_ = 1.409; p> 0.05) (table 2B, figure 2B).

The breeding female spent 54% of the time while she was recorded on camera traps, responding to stimuli, and 46% of that time she spent on other non-stimulus behavior. The female spent the most time when she was captured on recordings (32%) on sniffing stimuli (mean 23%, SE ± 0.087), then scratching the ground (16%, mean 12%, SE ± 0.059). The female spent the last time overmarking, it constituted 6% of the entire time during which she was visible on the recordings (mean 4%, SE ± 0.02). The time difference between the duration of sniffing unknown wolf and unknown dog stimuli is statistically significant (F = 3.612; p <0.05) however, no statistically significant results were obtained after the Tukey HSD test (table 2B, figure 2B). In the case of overmarking and scratching of the ground, the time differences between the reactions to unknown wolf and unknown dog urine are not statistically significant (respectively, *F*_2,36_ = 0.703; p> 0.05 and *F*_2,36_ = 1.022; p> 0.05).

Finally, we also checked how a single juvenile would react to stimuli. A single juvenile spent 157.2 seconds sniffing stimuli, which accounted for 51% of its activity (mean 46%, SE ± 0.05). The remaining 49% of the time of their activity, the juvenile devoted to other non-stimulus behaviors (mean 54%, SE ± 0.05). Unknown wolf stimuli were sniffed by a single juvenile longer than that unknown dog stimuli, moreover, the difference between an unknown wolf and an unknown dog is statistically significant (*F*_2,36_ = 5.974; p> 0.01) (table 2B, figure 2B).

## Discussion

Improved protection of carnivores in Europe creates a fascinating situation, when wolves and dogs increasingly share the same landscape and can potentially interact. This raises a lot of questions about the possibility of intermingling domesticated and wild environments, and the consequences of their interactions with each other. Some of these questions have already been approached, but most of them are still not answered. In our research, we showed that wolves in the natural environment showed a clear behavioral response toward dogs. This means that the presence of dogs in wolf territories does matter and therefore more emphasis should be placed on exploring the consequences of that situation.

Our study revealed that wolves do react to unknown dog smell. However, they reacted less intensively to the smell of an unknown dog than to the smell of an unknown wolf. The time the wolves were sniffing the stimuli from the unknown wolf was significantly longer than the wolves were the stimuli from the unknown dog, however, in the case of overmarking and scratching the ground, the difference between the response time to both stimulus was not significant. Wolves’ interest in the scent of an unknown wolf, expressed by long sniffing of the stimulus, may be related to the defense of their territory against potential intruder from the same species (Bekoff 2001), foreign wolves, while significantly shorter sniffing of the dogs’ stimuli may suggest that wolves do not associate the dog with a threat. However, in such a situation, one would expect wolves to similarly distinguish dog smell during overmarking and scratching the ground. Both behaviors are clearly related to the defense of the territory, and when performed together by the breeding pair, they are a strong signal of readiness to fight with the competition.

We hypothesize that wolves spend less time sniffing the stimulus from the unknown dog due to the frequent presence of dogs in the Kampinos National Park (Diserens et al., unpublished data). This may create a “scent pollution” effect, that has a disruptive effect on the natural behavior of wolves. In such a case, a similar experiment in a place with lower anthropopression and little presence of dogs in wolf territory should show no difference in the time that wolves would spend sniffing stimuli from the unknown dog unknown wolf stimuli.

Young wolves spend much more time exploring dog urine compared to adults. They also spend a similar amount of time sniffing both stimuli. The possible explanation for this behavior is that they do not yet distinguish between a dog smell and a wolf smell. They need to learn, and the question is – does a high abundance of dogs in Kampinos National Park, help them?

The interest of wolves in smells from other individuals of their species, as well as reacting to these smells by overmarking them, is a visual and chemical method of preventing aggression between individuals, thus the possibility of being injured or killed (Gosling 1982). However, the scent marking process itself is expensive, therefore wolves mark only the most important parts of their territory (Zub et al. 2003) and intensify marking in particularly sensitive periods, for example during heat (Peters and Mech, 1975; Zub et al. 2003) and care over pups (Llaneza et al. 2014). Moreover, wolves choose places where their markings can be easily located by other individuals (Barja et al. 2004; Stępniak et al. 2020). In this context, all actions performed by wolves in the KPN to protect their resources should result in intruders withdrawing from their territory. Therefore, the time spent by wolves reacting to dogs’ scent marking may be treated as an action not bringing the expected effect. Most dogs appearing in KPN, regardless of whether they are led on a leash or not, are under human control, which limits their natural behavior and is a major factor in determining where they will move. Thus, overmarking dogs’ scent markings by wolves cannot cause them to stop appearing. As a result, wolves spend valuable energy resources on behavior that has only a limited effect.

The average time spent on the reaction to the scent stimulus coming from the dog is 4.4 s in the case of a juvenile, and in the case of a breeding pair, it is 3.8 s and 3.14 s (female and male, respectively). During the study period, the dogs were recorded 91 times, which means that if each of these dogs appearing, at each visit, would leave only one scent marking, to which later passing wolves would react with a similar frequency as to the stimuli proposed in the above experiment, it is the juveniles that would spend an additional 160s, and the breeding pair an additional 80s and 89s (female and male, respectively) only responding to these stimuli. These calculations do not consider many factors, such as the fact that canines are able to distinguish odors from the same individuals, and therefore react to them less with each subsequent contact (Brown and Johnston, 1983). However, since those studies were conducted in captivity, it is difficult to assess whether such habituation would also be present in natural conditions. The hypothesis put forward by Gosling (1982) allows for the assumption that territory owners will react with a similar commitment to scent marking every time they come across it since their main goal is to remove a foreign smell from their territory.

The importance of the interactions between wolves and dogs based on scent marking has so far been underestimated. The most frequently discussed aspects of the interaction between these two predators are hybridization, parasite transmission, or competition for resources (Lascureux and Linnell, 2014), hence it is impossible to compare the results of this study with other studies in this thematic scope. However, a similar experiment on the Australian dingo (*Canis dingo*) showed that they reacted more intensely to the smell of domestic dogs than those of their own species, which is not in line with the main result of our study. However, the authors assumed that such a reaction was related to the fact that the tested dingo already knew the smell of other individuals of their own species, whose urine was used in the experiment (Van Bommel and Johnson, 2017). Research on the possibility of using predator scents in the protection of farm animals (biofencing) showed that the tested wolves in the second year of the experiment responded less to scent stimuli from unknown wolves.

The authors explained this by the fact that scent stimuli were placed in the wolf’s territory only for a short time, not maintaining continuity, which could mean that residents in the following year did not consider their presence as a threat, like in the first year (Ausband et al. 2013). However, in the case of the Kampinos National Park, where the presence of dogs lasts all year round, it does not seem that a similar mechanism would be possible.

Wolves living in the Kampinos National Park reacted to stimuli from an unknown wolf and from an unknown dog, which confirmed that territorial wolves react to smells from unfamiliar individuals (Peters and Mech 1975). The smell from an unknown individual is a source of important information about potential partners or a potential threat (Lisberg and Snowdon 2009). However, we did not expect the juveniles to respond for a longer time than their parents to both scent stimuli, these results can only be obtained through direct observation.

The existing knowledge assumed that mainly breeding individuals are responsible for defending the territory against intruders (Peters and Mech 1975), however, the latest research has shown that juveniles also take an active part in it by marking the area with their own scent (Stępniak et al. 2020). Our results suggest that juvenile wolves learn from their parents when traveling through the territory with them. Therefore, the presence of dogs in wolf territory may obstruct process of learning.

### Are dogs a threat for wolves?

The fact that wolves are actively recolonizing landscapes previously used mainly by humans and their dogs mean that interactions between wild and domesticated predators will become more frequent. We can assume that most of those encounters will not be direct, but our research indicates that there is a form of communication between wolves and dogs, and this communication can influence the behavior of wolves. Based only on the results of our research, it is impossible to unequivocally confirm whether wolves regard dog scent as a threat signal or an harmless distraction. Adult wolves do not spend a long time exploring “unknown dog” scents, which might suggest that they do not see them as a threat. On the other hand, they overmarked them with the same intensity as an “unknown wolf” scent, leaving a clear signal that the territory is already occupied by a group of wolves. Therefore, we need more research to answer the question of whether dog scent is a negative factor for wolf behavior.

At the end is worth to point, that regulations of the Kampinos National Park prohibit the introduction of dogs into areas under strict and active protection, with the exception of assistance dogs (order no. 15/2020 Director of the Kampinos National Park dated 31.07.2020 on the improvement of the Kampinos National Park). However, in study of Diserens (Diserens et al., unpublished data), dogs were the most abundant predator in Kampinos National Park, considering wolves, lynx, foxes, and badgers combined. If wolves living in that area need to react even for a small number of dogs’ markings, we can still assume that this might be more distracting than in a forest without so many dog visits.

